# Rapid and Accurate Interpretation of Clinical Exomes Using Phenoxome: a Computational Phenotype-driven Approach

**DOI:** 10.1101/275479

**Authors:** Chao Wu, Batsal Devkota, Xiaonan Zhao, Samuel W Baker, Rojeen Niazi, Kajia Cao, Michael A Gonzalez, Pushkala Jayaraman, Laura K Conlin, Bryan L Krock, Matthew A Deardorff, Nancy B Spinner, Ian D Krantz, Avni B Santani, Ahmad N Abou Tayoun, Mahdi Sarmady

## Abstract

Clinical exome sequencing (CES) has become the preferred diagnostic platform for complex pediatric disorders with suspected monogenic etiologies, solving up to 20%-50% of cases depending on indication. Despite rapid advancements in CES analysis, the major challenge still resides in identifying the casual variants among the thousands of variants detected during CES testing, and thus establishing a molecular diagnosis. To improve the clinical exome diagnostic efficiency, we developed Phenoxome, a robust phenotype-driven model that adopts a network-based approach to facilitate automated variant prioritization and subsequent classification. Phenoxome dissects the phenotypic manifestation of a patient in conjunction with their genomic profile to filter and then prioritize putative pathogenic variants. To validate our method, we have compiled a clinical cohort of 105 positive patient samples (i.e. at least one reported ‘pathogenic’ variant) that represent a wide range of genetic heterogeneity from The Children’s Hospital of Philadelphia. Our approach identifies the causative variants within the top 5, 10, or 25 candidates in more than 50%, 71%, or 88% of these patient samples respectively. Furthermore, we show that our method is optimized for clinical testing by yielding superior ranking of the pathogenic variants compared to current state-of-art methods. The web application of Phenoxome is available to the public at http://phenoxome.chop.edu/.

## Introduction

Individual Mendelian pediatric diseases are considered rare, yet an approximately 8% population worldwide are identified as having a genetic disorder before reaching adulthood^1^. Next generation sequencing (NGS) has rapidly changed the landscape of clinical genetics by enabling the researchers and physicians to make novel gene-disease associations^2^ and precise molecular diagnoses^3,4^. However, NGS-based clinical diagnostics remains challenging with only about 30% of the patients receiving a definitive diagnosis^5^. The diagnostic quest in a clinical exome test is complicated by the sheer volume of variants detected and the presentation of overlapping phenotypic characteristics in affected individuals^6^.

A carefully designed analysis paradigm is essential for high quality interpretation of a clinical exome test^7^. Clinical correlation, which includes concurrent analysis of the patient’s phenotype and genotype, is central to the overall clinical interpretation^8^. During this step, putative causative genes and variants contributing to the disease are identified. Nonetheless, clinical correlation is often time consuming and requires extensive expertise in both medical genetics and genomics^9,10^. To perform clinical interpretation at scale and keep up with the latest discoveries that would require re-analysis, it is critical to engage computational algorithms to automate this procedure.

Using prior biological and clinical knowledge, such as previously known disease genes and pathogenic mutations, together with phenotype information may assist in gene-disease clinical correlation^11^. A number of databases^12-14^ that curate gene-disease associations have been developed, along with several computational variant annotation tools^15-17^. Specifically, phenotypic information has been recognized to have greatly enhanced the diagnostic power, prompting numerous phenotype-driven approaches that often employ machine learning methods, including eXtasy^18^, Phenomizer^6^, PHIVE^19^, Phevor^20^, PhenIX^21^, Phen-Gen^22^, SimReg^23^ and Phenolyzer^24^. Most of these tools use the vocabulary from the Human Phenotype Ontology^25^ to describe a patient’s phenotypic abnormalities.

These tools have clearly demonstrated the utility of using gene-curated phenotype data to improve disease gene identification. Most of these approaches have been validated on large number of simulated scenarios but often limited numbers of clinical samples. However, these purely computational approaches have been shown less effective on clinical cases compared to clinician-aid strategies^26^. Additionally, none of these tools have been validated on a large-scale clinical sequencing cohort.

Here, we present a computational framework, Phenoxome, to filter and then prioritize candidate variants using population frequency, deleteriousness and clinical relevance of the affected gene (Figure 1). Phenoxome uses two inputs, (i) a variant call format (VCF) file representing the genotype of the affected individual, and (ii) a set of symptoms described using Human Phenotype Ontology (HPO) terms. Our approach first filters the variants according to rarity, predicted protein effects and other prior knowledge. Following this, Phenoxome generates a personalized gene panel (PGP) according to the phenotype manifestation, and each gene in the PGP is scored based on its involvement in these phenotypes. Finally, each variant is prioritized based upon a composite score combining the knowledge inferred from both variant level and gene level. In this work, we first evaluate the performance of our method on comprehensive computational simulations of different scenarios. We then demonstrate the effectiveness of Phenoxome using 105 positive clinical exomes at The Children’s Hospital of Philadelphia (CHOP). Our approach outperforms state-of-art methods by yielding superior rankings of the causative variants on the clinical samples.

**Figure 1.**
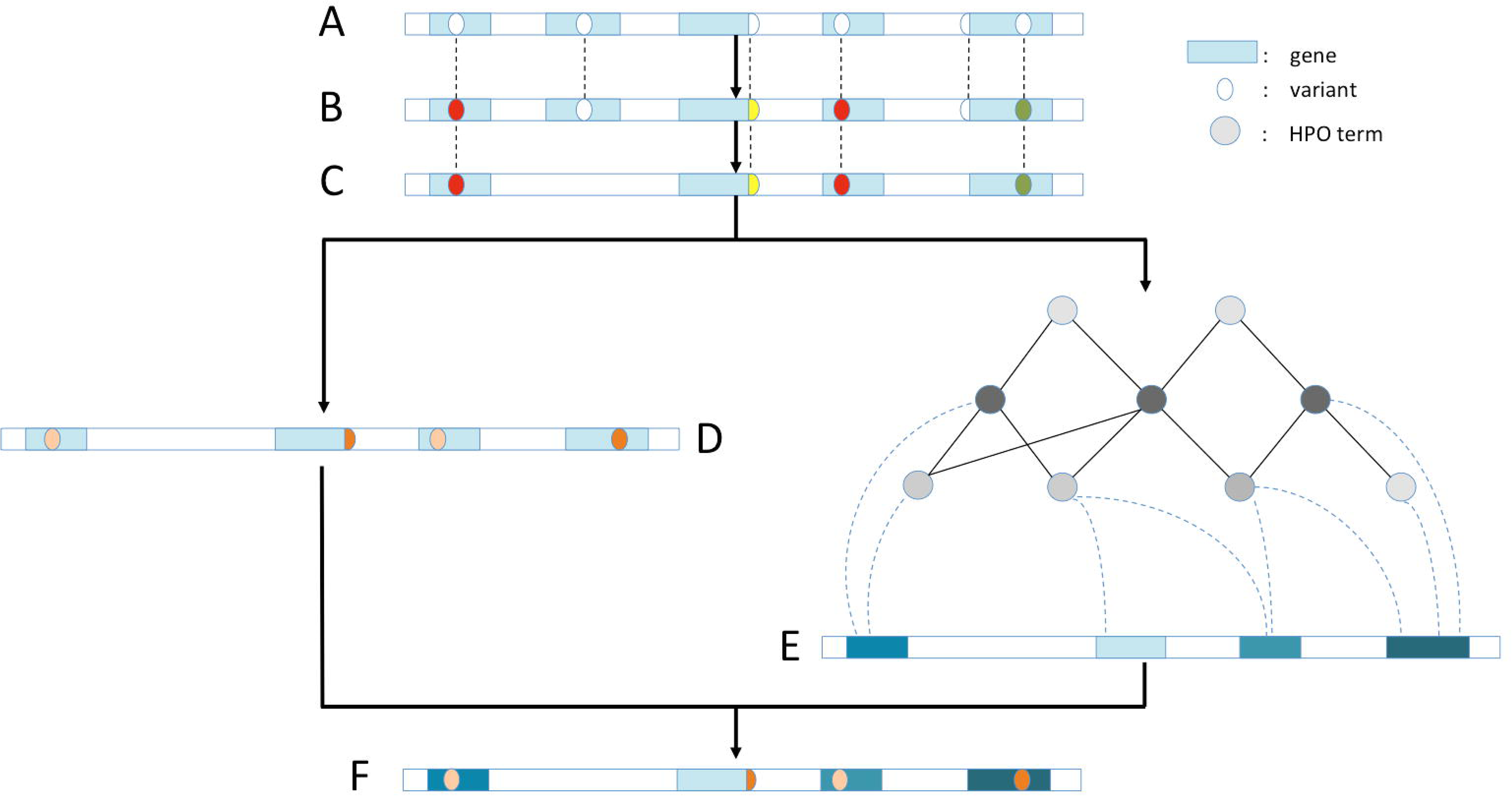
Overview of Phenoxome Workflow. A. Raw variants yielded from sequencing the patient’s exome and subsequent bioinformatics analysis. Blue rectangles imply genes and ovals indicate variants. B. Variants annotated by Phenoxome using a series of bioinformatics resources. Distinct color schemes indicate different predicted effects on protein products. C. Variants retained after filtering procedure depending on HGMD annotations, population allele frequency and functional effects. D. Variants deleterious score are derived from the tier strategy where a darker color implies a more disruptive variant. E. Genes harboring post-filtered variants are assigned phenotypic relevance scores inferred by their associations with relevant phenotypes in HPO. A darker color implies the gene is more pertinent to the patient’s phenotypic manifestation. F. Each of post-filtration variants receives an overall score by integrating both variant deleterious score and the gene’s phenotypic relevance score. Hence a global prioritization of the variants is achieved in the framework.

## Material and Methods

### Variant Annotation and Filtration

Our approach first annotates variants using SnpEff package v4.2^16^ with UCSC RefSeq/refGene database^27^. In addition, the variants are also annotated with a public version of the Human Gene Mutation Database (HGMD) database, obtained through Ensembl annotation system^28^ and minor allele frequencies from the Genome Aggregation Database (gnomAD) v2.0^29^.

Next, Phenoxome retains a variant if it meets one of the following criteria:

- AF < 1% in gnomAD and classified as disease mutation (DM or DM?) in HGMD
- AF < 0.2% in all sub-populations in gnomAD and predicted to alter protein or splice sites (i.e splice acceptor/donor, stop retained/gained, start/stop loss, inframe deletion/insertion, frameshift and missense variants)

Detailed variant filtering schemes are demonstrated in Figure 2.

**Figure 2.**
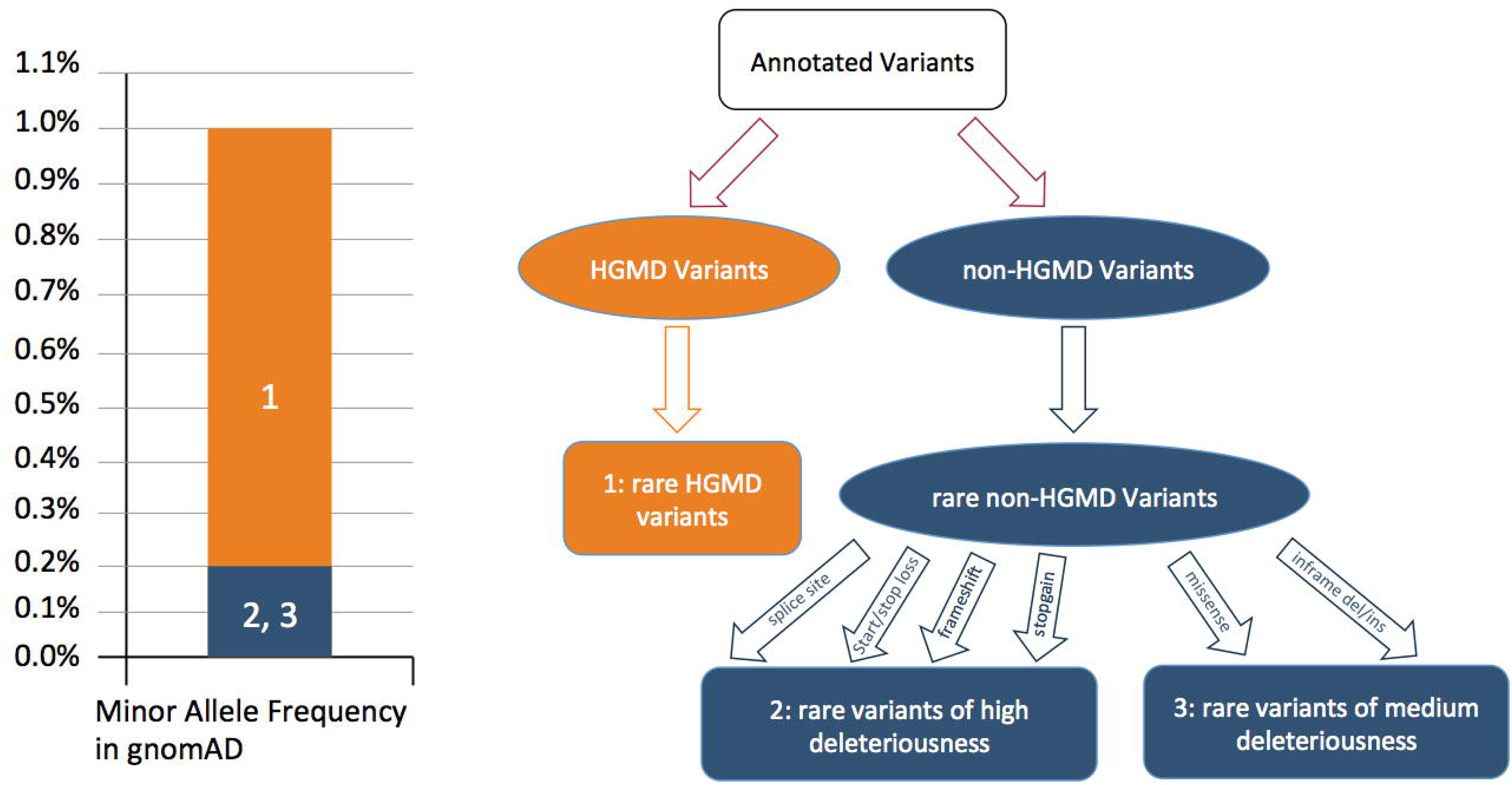
Variant Filtration Strategy and Tiers of Deleterious Score. A HGMD variant of DM or DM? class is retained if the minor allele frequency of the variant is less than 1% in general population in gnomAD database. This variant is binned tier one and is assigned 1.0 as deleterious score (shown in orange 1). A non-HGMD variant is retained if the predicted protein effect is disruptive, and its minor allele frequency is less than 0.2% in general population as well as five sub-populations (East Asian, Finnish, Non finnish European, African/African American and South Asian) in gnomAD. If the predicted effects of the variant include change of splice site, start/stop loss, frameshift and stopgain, the variant is binned tier two and is assigned 0.8 as deleterious score (shown in blue 2). The variant is binned tier three otherwise and is assigned 0.6 as deleterious score (shown in blue 3).

### Variant Prioritization Strategy

In Phenoxome, each variant that passes filtration receives a comprehensive score reflecting its likelihood of being pathogenic to the affected individual, and hence a global ranking of the variants is achieved based upon the scores. Similar to other approaches^21,22^, the composite score of each variant is contributed by deleterious score and phenotypic relevance score, derived from variant level and gene level respectively. A variant level score usually indicates the deleteriousness of the variant, inferred by characteristics such as rarity, evolutionary conservation and predicted functional impact^30,31^. In general, a gene level score reflects the assessment of the affected gene’s functional involvement in the observed phenotypes. Unlike other approaches that calculate the composite score by averaging the variant score and the gene relevance score^19,21^, our approach assigns greater weight to the phenotypic component while generating the overall score of a variant. This empirical implementation was derived from the clinical observation that most of the rare variants with disruptive protein effects were harbored by genes that shared little or no known disease overlap to the phenotypic manifestation of the affected individual.

### Deleterious Score

Each of the variant that passes filtration is evaluated and assigned a deleterious score based upon its rarity, HGMD label and predicted functional impact. Inspired by clinical protocols classifying variants into different categories^32,33^, we implement a tier system to triage the variants into three different bins. A variant is deemed the most damaging if it is in HGMD with a DM/DM? class (bin 1). The damage level is deemed high if the functional impact of a non-HGMD variant is any of the following: splice site aberration, frameshift, stop gain, start loss or stop loss (bin2). The rest of the variants are deemed medium damaging with predicted effects including missense, inframe deletion or inframe insertion (bin3). Binned variants are given a deleteriousness score of 1.0, 0.8 and 0.6 for bins 1,2 and 3 respectively (Figure 2).

### Phenotypic Relevance Score

Variants are also assessed on the gene level through the utility of HPO. HPO is a computational representation of a wide collection of phenotype abnormalities in human. Each of the phenotypes in the vocabulary is annotated with genes implicated with the clinical symptoms, curated from resources including OMIM^13^ and Orphanet^34^. Because of its strictly controlled and standardized vocabulary, hierarchical structure and well-defined phenotype-gene relationships, HPO has become an ideal resource for clinical phenotyping^35^.

Phenotypic terms in HPO are organized in a directed acyclic graph where they are associated by “is a” relationships. An “is a” relationship indicates that one phenotype is a subclass of another phenotype that is a more generic parent term^36^. For instance, Abnormality of the atrial septum “is an” Abnormality of the cardiac setpa which “is an” Abnormal heart morphology. The design of Phenoxome takes the advantage of the hierarchical structure of HPO and assembles a Personalized Gene Panel (PGP) for each patient, where each gene of the PGP is potentially associated with the input phenotypes. Our approach starts from each of the provided phenotypes, and then traverses down the ontology tree to retrieve all of its direct and indirect subclass nodes/phenotypes until a terminal branch is encountered. The nature of “is a” associations guarantees that all of the children nodes are essentially subclasses of the primary phenotypes by the definition of the hierarchy. In addition, in order to account for imprecision in selecting the primary phenotypes in clinical scenarios, the algorithm also visits the immediate parent nodes of the input phenotypes. The original terms describing the phenotypes of the patient are considered primary, while the terms retrieved during the extending process are termed secondary. The algorithm to generate secondary phenotypes is demonstrated in Figure 3. Following this, PGP is compiled to collect all of the genes associated with any of the primary or secondary phenotypes. These genes are reported to have caused corresponding symptoms and therefore are potentially relevant to explain the patient’s phenotypes.

**Figure 3.**
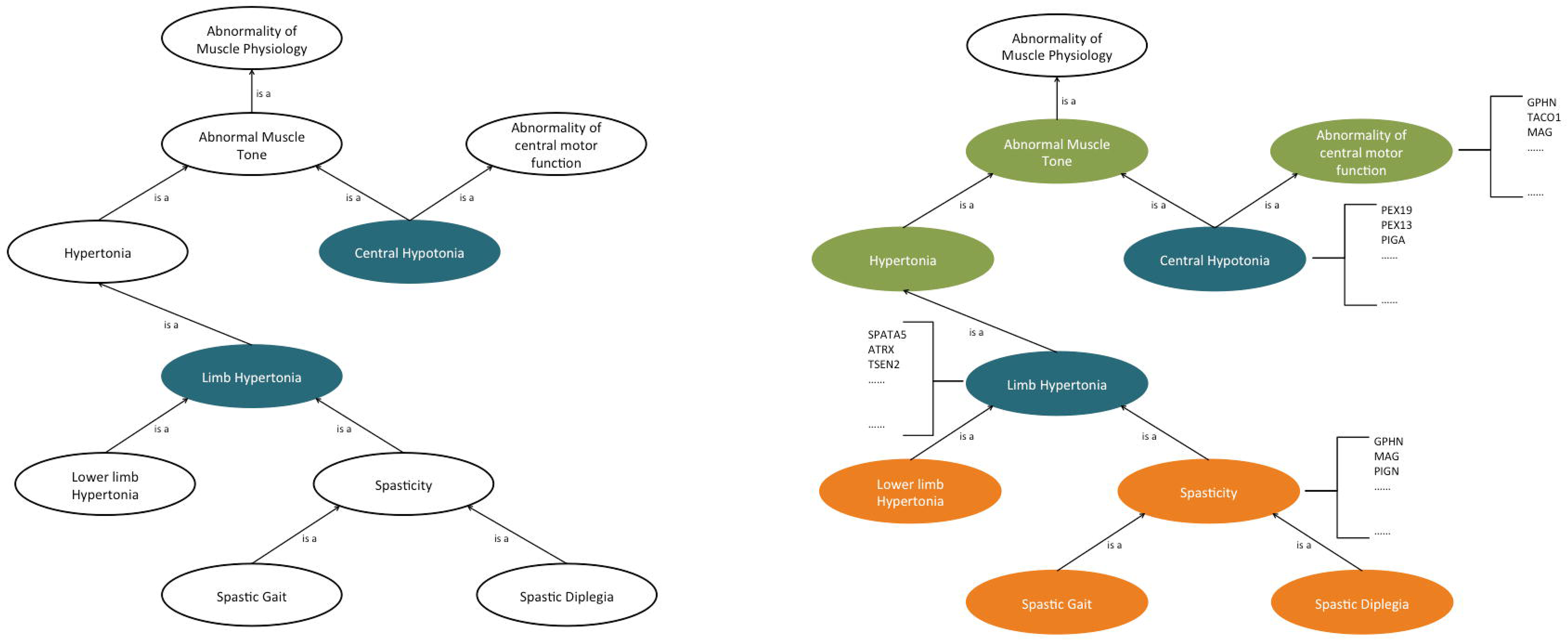
Compilation of Personalized Gene Panel using HPO. The cartoon panels demonstrate the procedure to generate personalized gene panel (PGP) from patient’s phenotypic manifestation using HPO in Phenoxome.The left panel is a representation of the initial state of a sub-network of HPO containing relevant phenotypes. Limb hypertonia and Central Hypotonia are the reported phenotypes in the patient example, which are the primary phenotypes and are highlighted in blue. The right panel reflects the end state of the sub-network where pertinent phenotypes are retrieved and personalized gene panel generated. The phenotypes in orange are children nodes of the primary phenotypes while the phenotypes in green are the immediate parents of the primary phenotypes. Genes associated with any of the primary or secondary phenotypes are compiled in PGP, such as GPHN (Spasticity and Abnormality of Central Motor Function) and SPATA5 (Limb Hypertonia).

Once Phenoxome identifies the primary and secondary terms, a sub-graph of HPO tree containing all of the nodes and their parent-child relationships is also generated (see Fig 3), we then employ a network-based stochastic approach, PageRank with Priors^37,38^, to prioritize each phenotype in the sub-graph. The algorithm evaluates the significance of each node of a graph with a clear-defined transition matrix by imitating a random walker surfing the graph. Starting from a root node, the surfer selects an outgoing edge from the current node randomly to jump to the next node in each iteration. The algorithm converges when the significance scores of the nodes become steady. In a directed acyclic graph, this process is similar to the ontology propagation described by Singleton et al^20^. However, with a set of priors (root set), the random surfer opts to jump back to any of the node in the root set regardless of its current location with a predefined probability in each iteration. The iterative stationary probability equation of a node n is given by

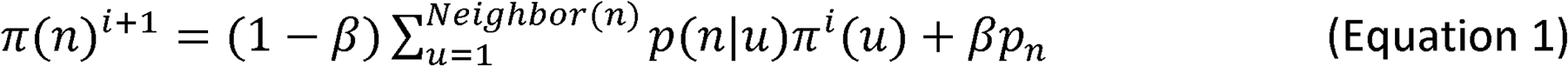

where *β* is the back probability. The first component of the equation summarizes the likelihood of arriving at this node from all of its neighboring nodes while the second component indicates transporting back to the root set. In Equation 1, *p*_n_is a 0/1 prior vector where 1 indicates the node is in the root set and 0 indicates otherwise. The stationary distribution after the convergence of the algorithm represents the probability of the random surfer landing on each node at any given moment.

In order to implement the algorithm in the context of the sub-graph of the HPO tree, we set the primary HPO terms as the priors and the back probability *β* to be 0.5 as it was suggested to yield optimal performance by previous studies^39^, meaning there is a 50% chance of the random surfer returning to the primary terms in each step. It is intuitive to see several benefits with this implementation to prioritize the phenotypes for the clinical utility. The primary phenotypes are ranked high because of the back probability; the secondary phenotypes that are close to the primary phenotypes are ranked high because they are easily accessible from the root set; and the secondary phenotypes with more “is a” relationships are ranked high because they are more likely to be visited during the random walk. Each gene in PGP may be associated with multiple primary and secondary phenotypes, thus a variant receives a phenotypic score that is the sum of the phenotypes’ scores the affected gene is associated with. In this way, variants harbored by genes associated with more significant phenotypes are ranked higher.

### Integrated Variant Pathogenic Score

As discussed earlier, an overall score is assigned to each of the candidate variants. The first component is the gene-phenotypic score and the second is the deleterious score. Weight factor *α* is employed to combines the two components together in the final significance score:

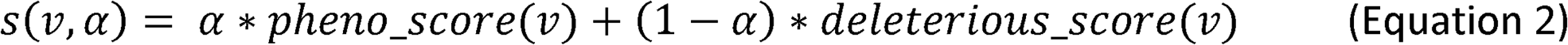

where *α* is intuitively set to 0.8 to ensure the global implementation is phenotype-driven. Thus, a prioritization of the variants is achieved based upon the final scores.

### Clinical Sample Cohort

Acquiring validated large-scale clinical cohorts for differential clinical diagnostics studies is challenging for a number of reasons. Thus, limited access to these resources has forced most of the abovementioned computational tools to perform their validation largely or solely using *in silico* patient profiles. We have collected a large cohort of clinical patients (n=105) where each individual patient received a positive molecular diagnosis from Clinical exome sequencing (CES) analysis. Eighty-five positive samples were from patients diagnosed by the clinical exome test at the Division of Genomic Diagnostics (DGD) at CHOP between 2014 and 2017. Twenty positive exomes were collected through the CHOP Pediatric Sequencing (PediSeq) project, which was a part of the National Human Genome Research Institute (NHGRI) Clinical Sequencing Exploratory Research (CSER) consortium. For this study, we define positive cases as having at least one pathogenic variant in the final clinical lab report. These pathogenic variants were thoroughly evaluated and classified as disease causing with concrete supporting evidence that was previously established such as published articles and clinical reports. They were confirmed by an orthogonal technology (Sanger Sequencing) before being reported. All of these positive cases had gone through comprehensive manual clinical correlation performed by experienced clinical geneticists during variant interpretation and the findings were confirmed by clinical laboratory directors.

The phenotypic features of these patients were carefully discussed and documented by physicians upon clinical chart reviews, and the corresponding HPO terms were selected to best represent the symptoms. A total of 422 unique HPO terms were used to describe the phenotypes of all individuals in the cohort. The numbers of HPO terms per case range from 1 to 23, with an average of 6 terms. Out of the 422 phenotypes, 306 terms (73%) were unique to just one individual respectively while another 62 terms (15%) were shared by only two individuals. This collection of phenotypes represents a wide range of developmental abnormalities. The most frequently used 15 terms are shown in Figure 1S. It should be noted that the phenotypes with the highest term frequencies are more generic than those with lower term frequencies. For instance, *Global developmental delay* was used to describe 43 patients in the cohort and the term was annotated with 1063 genes, Short stature was observed in 19 patients and the term was annotated with 832 genes. It was similar for terms such as *Failure to thrive* (16 patients, 577 genes), *Microcephaly* (14 patients, 678 genes) and etc. In contrast, terms that were unique to one or two patients were likely to be more specific, such as *Prominent fingertip pads* (1 patient, 8 genes), *Vocal cord paralysis* (1 patient, 11 genes) and *Metatarsus adductus* (2 patients, 30 genes).

### Sequencing and bioinformatics pipeline

For each of the clinical CES samples from the DGD, Agilent SureSelect Clinical Research Exome (CRE) V1 capture was used as the exome capture platform. FASTQ data generated by MiSeq/HiSeq was aligned to the hg19/Grch37 reference genome using Novoalign from Novocraft (Selangor, Malaysia). Read alignment and variant calling were performed with an in-house bioinformatics pipeline that incorporated Novoalign for read alignment, Picard (Broad Institute, Cambridge, MA) for marking duplicates, the Genome Analysis Toolkit (Broad Institute, Cambridge, MA) for variant calling (reference sequence: hg19/Grch37 build) and variant read depth filtering (≥5X). Variant call sets were generated in the format VCF 4.1/4.2. Variant annotation and initial variant filtration were then performed. This filtration restricted the data to variants in the Human Genome Mutation Database (HGMD) and/or rare variants with a coding effect such missense, stop loss, stop gain, start loss, in-frame insertion, in-frame deletion, frameshift insertion/deletion, and variants within the consensus splice site (+/-6 bases in intron) among all genes. To eliminate lab-specific sequencing artifacts we used an internal cohort compiled from 665 unaffected and unrelated clinical exome sequencing samples in the laboratory of DGD. Patient variants present at a high frequency (1.0% or > 6 individuals) in this cohort were removed. All 85 DGD clinical exome samples were filtered against this internal cohort. The VCF files of final products of these exome samples were generated as the input to Phenoxome.

For PediSeq samples, exome capture was performed using SureSelect V4 kit (51MB target size). Exome sequencing was done at 100X depth of coverage on HiSeq 200 sequencers. The same bioinformatics processing was performed on these samples and the variant call sets were filtered to remove common variants using the Exome Aggregation Consortium (ExAC)^29^ database (>0.5% MAF) and the “internal cohort (>5% MAF)” to remove high frequency technical artifacts. The internal cohort consisted of data from 269 samples from various PediSeq cohorts.

## Results

### Ranking Candidate Genes Using Synthetic Patient Profiles

Since the phenotypic scores of candidate variants are imperative to the overall prioritization and due to the general lack of clinical data, we first assessed the performance of the candidate gene ranking through *in silico* patients^6,23,24^. We focused on 33 monogenic diseases with known causative genes and used a similar strategy discussed by Masino et al^40^.

Each of these diseases is well characterized with sufficient phenotypic features and penetrance information as described using HPO terms. We produced synthetic patient profiles by selecting disease HPO terms for the patient, with the probability of being affected determined by the penetrance data. For instance, to generate a synthetic patient profile with a disease of phenotypes A that is observed in 70% of the patients with this disease, a random value between 0 and 1 was generated by a random number generator of uniform distribution. If the number was equal to or less than 0.7, then phenotype A was assigned to the profile. Phenotype A was not added to the synthetic profile otherwise. Same procedure was carried out on other phenotypes associated with this disease. This process was repeated to generated 1,000 synthetic profiles of each disease. This scenario was denoted as the “optimal” clinical scenario where the patient phenotype is well recognized.

However, it is common that patients may exhibit symptoms that are not directly related to their indication for testing. In addition, clinician or medical specialists may fail to choose the most accurate terms to describe the phenotype manifestation or the clinical anomaly is not properly defined in HPO vocabulary, so a more generic phenotype is used. These two typical scenarios contribute to a large number of cases in clinical practice and are denoted as “noisy” and “imprecise” respectively. In order to imitate the “noisy” scenario, randomly selected HPO terms were added to the “optimal” profile as noise. The number of noise terms was determined to be half of the number of “optimal” terms in the synthetic patient. To investigate the impact of the “imprecise” terms, each of the “optimal” terms of a patient was replaced by a parent term of the original HPO term ^40^. These procedures were repeated 1,000 times so that 1,000 *in silico* patient profiles for “noisy” and “imprecise” scenarios were generated respectively, resulting a total of 99,000 simulated profiles for the study.

We carried out candidate gene prioritization approach of Phenoxome on all simulated patient profiles of the three scenarios. For each synthetic patient, our algorithm first generated the PGP from the phenotypes and then prioritized the genes in PGP using the phenotypic relevance scores presented in Methods. In all 99,000 simulated cases, the causative genes were constantly captured by the PGP across the 33 diseases of the above scenarios. In the “optimal” scenario, the causative gene was ranked first for 98.5% of the cases. This was not surprising since the causative disease gene was associated with at least one phenotype of greater than 50% penetrance in 32 diseases except for *Holt-Oram syndrome*. For this disorder, which is caused by mutations in the TBX5 gene, the most common phenotypes were Defect in the atrial septum and *Hypoplasia of the radius*, with 41.46% and 37.8% penetrance respectively (Supplement file). Introducing the “noise” terms did not have any substantial impact on the rankings. In the “noisy” scenario, the causative gene was ranked first for 94.4% of the cases. Consistent with previous studies, a deteriorated performance of Phenoxome in the “imprecise” scenarios was observed where the causative gene was ranked first in only 3.7% of the cases. However, the target gene was ranked within top 10% of the PGPs in 89.8% of the cases. The overall summary of the performance of Phenoxome in three scenarios is demonstrated in Figure 4. The respective AUC (area under curve) values for the “optimal”, “noisy” and “imprecise” scenarios are 0.995, 0.991 and 0.952, which demonstrated a modest advantage compared to other network-based methods^6,40^.

**Figure 4.**
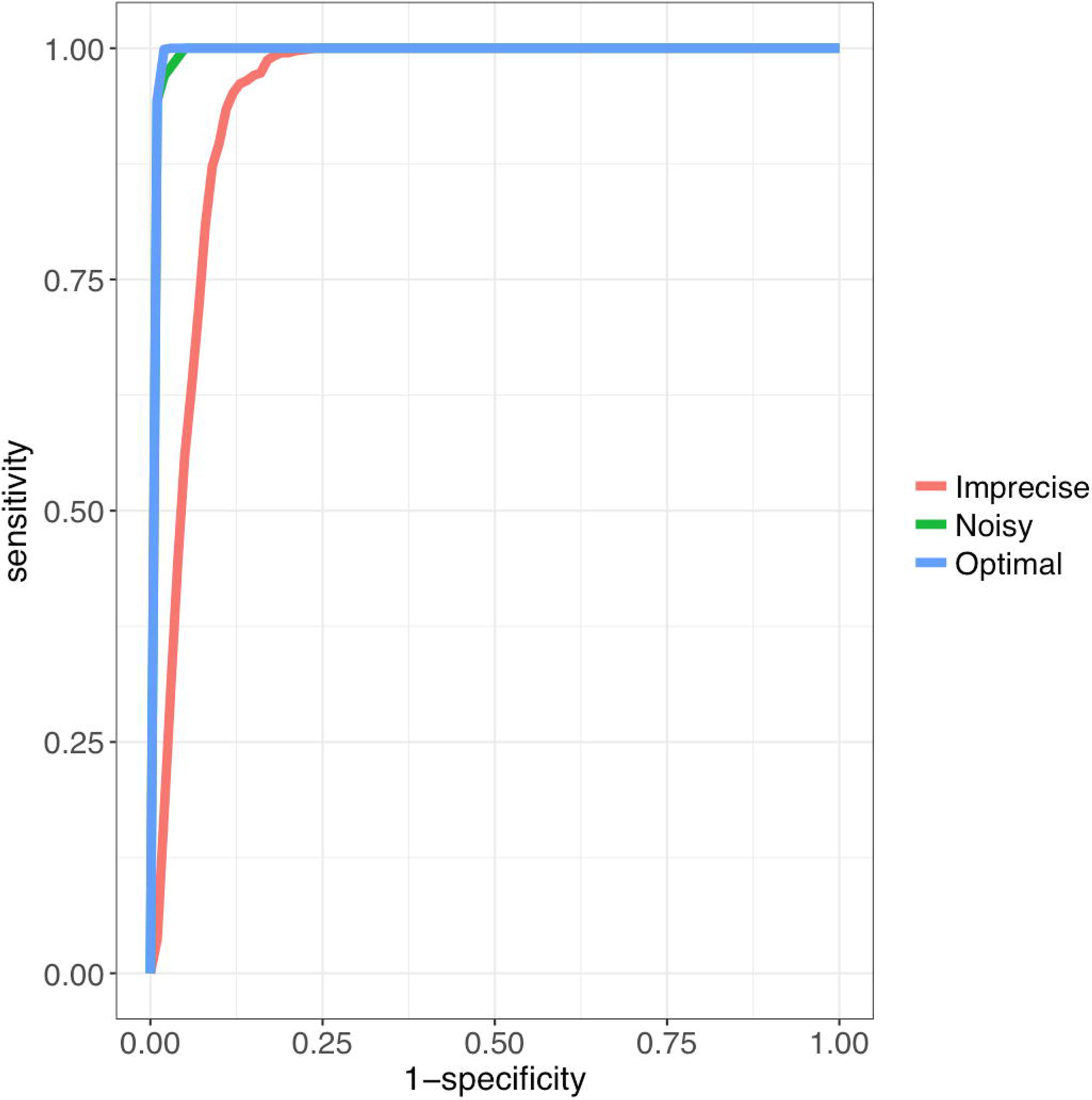
Performance of Phenoxome on Simulated Profiles. Blue curve is the ROC of Phenoxome’s performance on the “optimalscenarios” based on score ranks; green and red curves indicate the “noisy” and “imprecise” scenarios respectively. Each of the plots was generated from 33,000 simulated cases. Sensitivity was defined as the frequency of “target” genes that are ranked above a particular threshold position, and specificity as the percentage of genes ranked below the threshold. For instance, a sensitivity/ specificity value of 70/90 indicates that the disease gene (the “target”) is ranked among the best-scoring 10% of genes in 70% of the prioritizations.

### Performance on Clinical Samples

We then used the clinical cohort (n=105) to validate Phenoxome performance. Paired VCF files and HPO terms of each clinical sample were utilized as described in Methods. Phenoxome generated a list of ranked variants independent from the original clinical evaluation. The numbers of variants in the original input VCF files ranged from 37,150 to 258,968. Number of variants was reduced (431 to 2,995) after the effective variant filtration strategy implemented by Phenoxome. The clinically reported pathogenic variants were consistently reported in the final ranked lists and its rank was recorded for each of the patient. If more than one pathogenic variant was reported, the best rank of these variants was used in the benchmarking for the case.

The median rank of the pathogenic variants in the 105 patients was 5 with a standard deviation of 23. Specifically, 92 (88%) pathogenic variants were ranked in the top 25, 74 (71%) in the top 10, 53 (51%) within the top 5, while 17 pathogenic variants (16%) were ranked in the first place.

Causal genes harboring the pathogenic variants were captured in the PGP in 95 out of 105 cases (associated with at least one primary or secondary phenotype). The median rank of the pathogenic variants was 4 among those 95 cases. Three primary and/or secondary phenotypes were annotated to the causal gene on average among these cases. Causal genes were associated with at least one primary phenotype in 78 out of 95 cases while they were associated with only secondary phenotypes in the other 17 cases. No statistical differences between the two groups were observed regarding the final ranks of the pathogenic variants. For the 17 cases where the causal genes were associated with only secondary phenotypes, the ranking of causative variants were in the range of 1 to 24 with an average of 6. It is perennial that the causal gene was associated with a primary phenotype when it was also implicated in secondary phenotypes inferred from other primary phenotypes. For instance, a missense variant in *SCN1A* was reported as pathogenic for a patient from the cohort. *SCN1A* was not only associated with one of the primary phenotypes (Global developmental delay) but also annotated with *Obtundation* status and *Seizures*, which were secondary phenotypes inferred from the primary phenotype *Status epilepticus*. Hence, *SCN1A* was compiled in the PGP and the pathogenic variant was ranked in the first place by Phenoxome. In a more striking instance where the patient was documented with phenotypes of *Chronic mucocutaneous candidiasis, Recurrent fungal infections, Recurrent candida infections* and *Impaired T cell function*, the pathogenic variant was identified in *IL12RB1* that was not directly annotated with any of the primary phenotypes but was associated with *Onychomycosis* which was a sub-class of *Recurrent fungal infections*. Thus, the causal gene was captured in the PGP and the missense pathogenic variant was ranked in the second position for this patient.

Phenoxome heavily relies on the provided phenotypes and the gene-phenotype associations in prioritizing the variants. Thus, using the most accurate and up-to-date phenotypes is essential to achieving the optimal performance. On the other hand, as phenotypic features of patients evolve over time, as well as new gene-phenotype associates are uncovered, re-analysis using Phenoxome could yield new diagnosis.

In our clinical validation cohort, 10 pathogenic variants (marked in orange in Figure 5) were not in PGP during the initial benchmarking. The pathogenic variants in these 10 cases were ranked in the range of 5 to 140. With one exception where the pathogenic variant was ranked in top 5, the rest of these variants were all scored below the median rank of the cohort, with an average rank of 59. To investigate the ten cases, re-analysis was performed using the latest build of HPO (build 1249, January 2018). Three out of the ten causative genes were identified to be annotated with at least one pertinent phenotype in the re-analysis, resulting substantially better ranks of the pathogenic variants (Table 1). This improvement was due to novel gene-phenotype relationships curated by HPO that were absent in the HPO database version at the time of the initial analysis. These findings highlight the clinical utility of re-analysis of exome data to yield additional diagnosis in a systematic manner^41^. For the remaining seven cases, we noted that precise HPO terms were not provided in the clinical HPO phenotyping information. For example, in a case where the pathogenic variant was confirmed in *CTSA, Decreased beta-galactosidase activity* was mentioned in the clinical chart review, but was not provided to Phenoxome as a primary phenotype of the patient. With only two associated genes (*CTSA* and *GLB1*), this specific HPO term could have guided Phenoxome to include *CTSA* in the PGP. Instead, *Abnormality of lysosomal metabolism* was reported which was a more generic term, for it was more inclusive as the patient manifested other symptoms related to lysosomal metabolism. However, *Abnormality of lysosomal metabolism* resides in a separate trunk in the HPO tree compared to *Decreased beta-galactosidase activity* and hence was never visited or retrieved by the algorithm. Therefore, *CTSA* was not in the PGP and the overall rank of the pathogenic variant was poor.

**Table 1.** Re-analysis Results on Three Clinical Samples

| Sample | HPO terms | Initial analysis Rank | Re-analysis Rank |
| --- | --- | --- | --- |
| CHOP-PA-S28 | HP:0000577;HP:0001249;HP:0000717;HP:0000739;HP:0000047;HP:0004322;HP:0001999;HP:0002275;HP:0011098 | 41 | 31 |
| CHOP-PA-S52 | HP:0001263;HP:0001508;HP:0008872;HP:0001385;HP:0004474;HP:0004482;HP:0000280;HP:0010813 | 48 | 8 |
| CHOP-PA-S78 | HP:0000410;HP:0001508;HP:0001263;HP:0001738;HP:0000010;HP:0012330;HP:0100785;HP:0004322;HP:0000076 | 41 | 3 |

**Figure 5.**
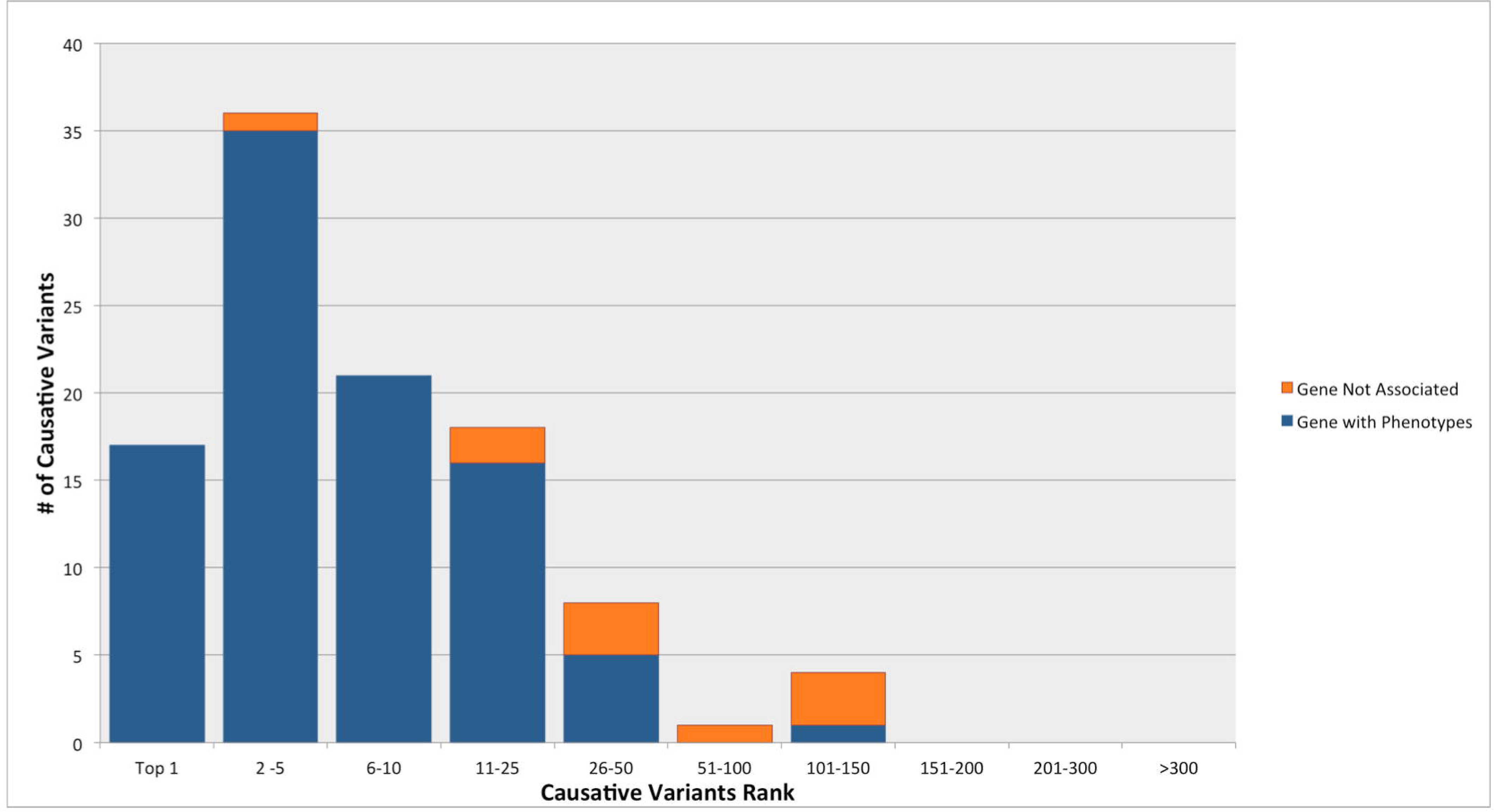
Performance of Phenoxome on Clinical Samples. Blue bars represent the cases where the causative variants/genes are associated with at least one pertinent phenotype. Orange bars represent the cases where the causative variants/genes are not associated with any pertinent phenotype. 53(51%) target variants were ranked in top 5 while 92(88%) target variants were ranked in top 25 among 105 clinical cases.

### Comparison to Other Methods

Unlike Phenoxome, most previously published computational approaches were primarily assessed using simulated patient data (see Table S1). It has been suggested that the performance of such tools, including Phenoxome, could vary significantly when using actual clinical cases^42,43^. Therefore, we compared the performance of Phenoxome against other methods using our clinical cohort. A recent comparative study examined the performance of a wide range of phenotype-driven variant prioritization methods, including OMIM Explorer^44^, Phen-Gen, Phevor and PhenIX, on 21 positive clinical exomes, and determined that PhenIX was the most effective^45^. Thus, we benchmarked the performance of PhenIX on the exomes in our cohort and compared the rank positions of the causative variants with Phenoxome.

The samples were processed through a local implementation of PhenIX as we could not use the web version to analyze the samples directly due to the clinical nature of the data. The latest version of the Exomiser (V8.0.0)^46^ was obtained through its ftp site and locally installed. To be consistent with the analysis on Phenoxome, same sets of paired VCF files and HPO terms were analyzed in the template of the local installation, where the variant prioritization method was set to PhenIX. A similar variant filtration strategy was implemented in the template to ensure a fair comparison. Variants of MAF greater than 0.5% that were annotated with functional impacts of intergenic, intronic, upstream/downstream gene or synonymous mutation were removed. PhenIX used the template as its input parameter to generate the ranking of the variants from the input VCF file specified.

As the result, the pathogenic variants were ranked in the range of 1 to 497 among 86 patients in the cohort. PhenIX ranked the causative variants in the first place in 22 (21%) out of the 86 cases, showing a slight advantage over Phenoxome (17/105). However, PhenIX scored notably fwer causative variants in the top 5 (45, 43%), top 10 (50, 48%) and top 25 (57, 54%) than Phenoxome respectively (Table 2). Moreover, PhenIX did not report the causative variants in the final ranked list in 19 out of the 105 cases.

**Table 2.** Performance Comparison Between Phenoxome and Phenix on **Clinical** Cohort

| 105 Clinical Cohort | Phenoxome | PhenIX |
| --- | --- | --- |
| Top Hit | 17 (16%) | **22 (21%)** |
| Top 3 | **41 (39%)** | 39 (37%) |
| Top 5 | **53 (51%)** | 45 (43%) |
| Top 10 | **74 (71%)** | 50 (48%) |
| Top 25 | **92 (88%)** | 57 (54%) |
| Top 50 | **100 (95%)** | 63 (60%) |
| 50 - 100 | **1 (1%)** | 9 (9%) |
| 101-150 | **4 (4%)** | 8 (8%) |
| >150 | **0** | 6 (6%) |
| Miss | **0** | 19 (18%) |

The statistical analysis on the overall rank positions of the pathogenic variants in the large clinical cohort between the two approaches indicated the superiority of Phenoxome over PhenIX (*p* = 0.0015; Mann-Whitney test). Collectively, Phenoxome outperformed PhenIX by yielding more robust rank positions of the pathogenic variants (Figure 6). Although PhenIX had several more pathogenic variants ranked in the first place, Phenoxome placed more pathogenic variants in top 5, top 10 and top 25 of this large cohort of genetic heterogeneity, and it did not miss the pathogenic variant in any of the cases.

**Figure 6.**
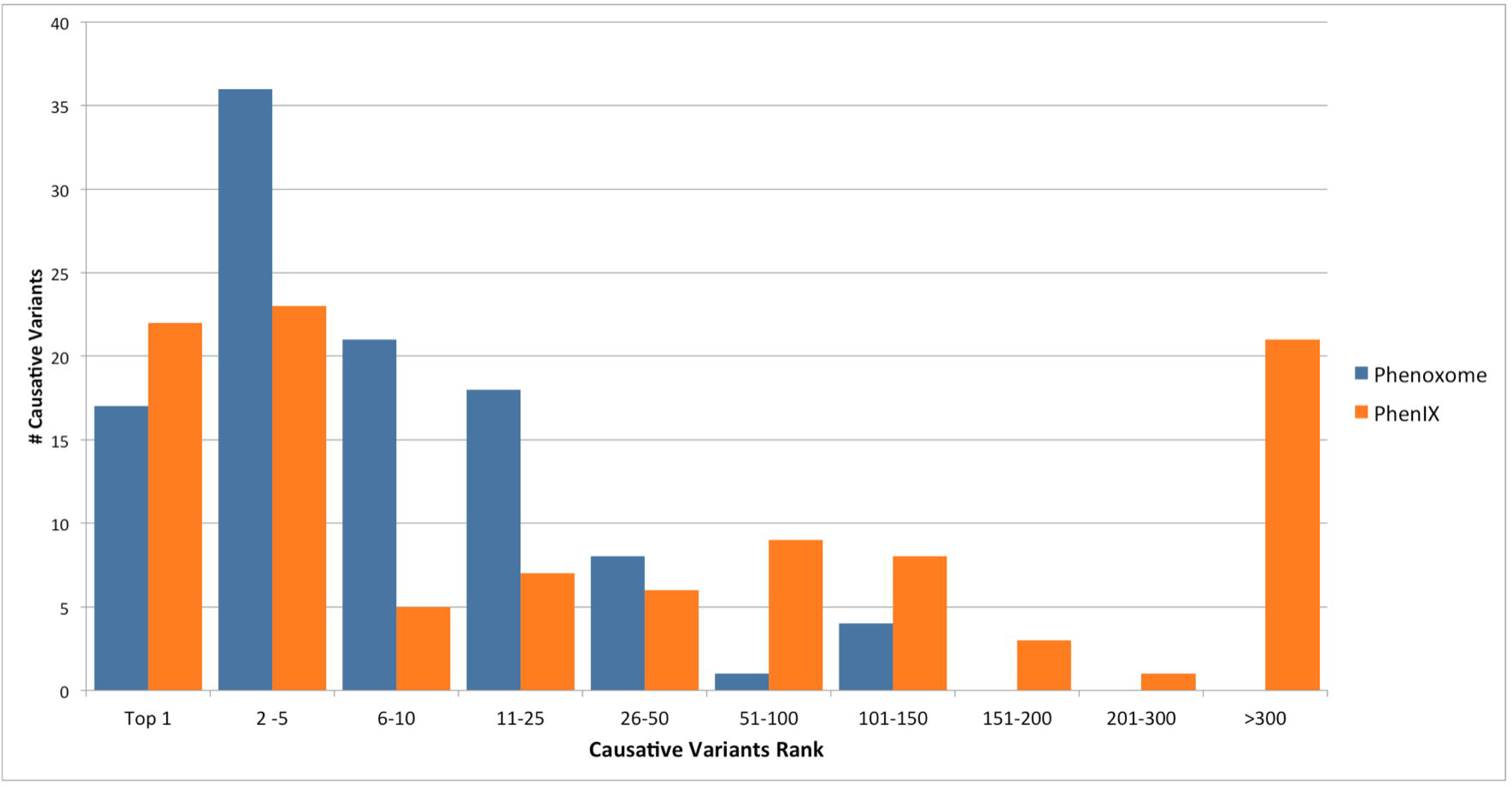
Comparison Between Phenoxome and Phenix on Clinical Cohort. Blue bars represent the rank of the causative variants in Phenoxome while orange bars represent those in Phenix. Phenix placed 22 causative variants on top among 105 cases, which was more than that of Phenoxome. However, Phenoxome ranked more causative variants in top 5, 10 and 25, demonstrating a superior performance overall. In addition, 19 causative variants were missed by Phenix (in >300 bar).

### Composite Score Weight Factor Revisit

The weight factor *α* of Equation 2 was used to combine phenotypic relevance score and the variant deleteriousness score to generate an overall score for each variant, and was set to 0.8 empirically to calculate the scores and hence prioritize the variants. After benchmarking Phenoxome’s performance on 105 positive clinical exomes, we re-estimated *α* to achieve an optimal result. In practice, the higher score of the pathogenic variant generated by Equation 2, the better the rank in each case. Thus, optimizing the global ranking is correlated to maximizing the final scores of all of the pathogenic variants:

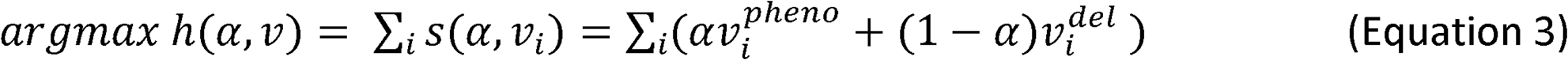

where *s*(*α, x*_*i*_) is Equation 2 for each case i in the cohort, *v*^*pheno*^ is the phenotypic relevance score of the pathogenic variant, *v*^*del*^ is the variant deleteriousness score and *α* ∈ (0,1). Then,

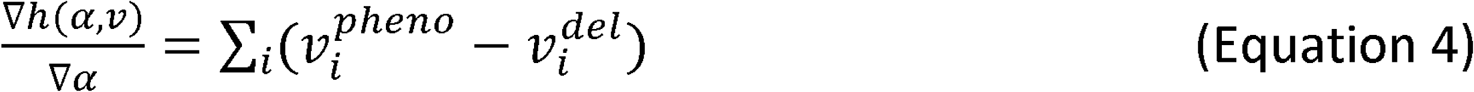

The gradient suggests that *h*(*α, v*) is maximized when *α* is close to 1. In practice, little gain was observed in the 95 clinical samples where the pathogenic genes were in the PGP when a was set to 1. However, completely eliminating the variant deleterious score from Equation 2 will substantially deteriorate the ranking performance of Phenoxome on those ten pathogenic variants whose genes were not in PGP. Thus, to reduce overfitting the model with our cohort, we decide to continue to use *α* = 0.8 to balance the two components in Equation 2 for the consistency of the variants prioritization in Phenoxome.

### Phenoxome Web Application

The methods described in this paper have been implemented in a web-accessible system using Java that runs on a Linux server utilizing an Apache Tomcat Server environment. Provided with paired VCF file and a list of HPO terms describing the phenotypes of an affected individual, Phenoxome performs the data analysis and then generates a report of ranked variants for download. The database backend of Phenoxome is updated on a monthly basis, and the web application is publicly available at http://phenoxome.chop.edu. To allow Phenoxome to run in a reasonable time frame (∼4 to 10 min) and with a limited memory, we limit the size of input VCF file to 10 MB.

## Discussion

We have presented a computational framework that is optimized for identifying causative variants utilizing genotype and phenotype information in a clinical context. Through a series of benchmarking using both *in silico* and large-scale exome clinical data, Phenoxome has demonstrated high clinical utility in identifying the causative variants in a wide range of scenarios. Further, our approach has outperformed current algorithms in the large cohort of heterogeneous clinical exome samples, by ranking most pathogenic variants in top 25 and in over 70% of the cases in top 10.

Phenoxome’s advantage over PhenIX is exhibited through the capability to consistently retain the causative variants during the filtration process and yield better rank positions overall. Specifically, we believe that our model outperforms the semantic similarity-based PhenIX in the clinical setting because Phenoxome is more patient-centric through utilization of PGP. In semantic similarity-based models, the phenotypic relevance score of a gene is calculated comparing the set of phenotypes manifested by the patient and all of the phenotypes associated with the gene, which may lead to what we call “phenotype dilution”. In clinical practice, the symptom manifestations of a patient are usually consolidated on several key phenotypes. However, a well-studied gene may be associated to a wide range of diseases that may be unrelated to each other. All of these associated phenotypes contribute to the semantic similarity calculation, which may “dilute” the associated phenotypes specific to this patient. On the other hand, our approach only takes into account the primary and secondary phenotypes of the patient and thus the signal is enhanced for the causal gene, as other irrelevant phenotypes associated to the gene are not considered in the analysis. For instance, in an exome sample where a patient was documented with *Volvulus, Intestinal pseudo-obstruction, Cholestatis* and *Intestinal malrotation*, a missense variant in *ACTG2* was classified pathogenic. *ACTG2* was associated with a total of 34 different phenotypes in HPO, ranging from *Camptodactyly of finger to Sepsis*, including *Intestinal malrotation*. Most of these phenotypes were not observed and unrelated to this patient, as they were “noise” in the similarity metrics and PhenIX prioritized this variant at rank 70. In contrast, Phenoxome did not consider those “noise” phenotypes associated with *ACTG2* in its modeling and ranked the causative variant in the second place.

Phenoxome does not make inferences from non-human genomic data, inlike several other tools^20,22^. By utilizing only well-established evidence of human disease and associated genes, it is not designed to make novel gene discoveries but rather is optimized for clinical testing. While this strategy restrains Phenoxome from leveraging information from other heterogeneous resources of genetic data, it offers clinical robustness that precludes non-human genomic data, which often does not offer validity to benefit clinical diagnostics^45^.

Another advantage of Phenoxome is its ability to perform very well with or without inheritance information. Trio-based clinical exome sequencing (both parents and their affected child sequenced simultaneously) has shown more effective in detecting de novo and compound heterozygous variants compared to proband-only approach^5,47^. However, parents are not always available for exome analysis. While Phenoxome can certainly rank variants prioritized by any inheritance model, it is also optimized to process and analyze singletons only and is currently agnostic to gene inheritance patterns.

It can be seen from the performance benchmarking that the computational tools behave differently on the clinical samples as opposed to synthetic patients. This highlights the importance of the validation of computational tools on real clinical data. Furthermore, our validation results also indicate that selecting the most accurate phenotypes to describe the symptom manifestation of a patient is crucial to a successful clinical diagnostic. The rankings of the pathogenic variants where the causal genes were captured in the PGP were consistently better than those that were not. We have shown that several scenarios that are not mutually exclusive could have contributed to those cases. One plausible explanation is the phenotype-gene annotations are absent in the database because HPO has missed them. The HPO project is partly dependent on text-mining algorithms to consolidate phenotypes and curate phenotype-gene associations from resource such as OMIM and PubMed Corpus^25^, as well as focused efforts to manually curate terms. However, it might not be sufficient to parse and extrapolate recently published articles or discoveries. In addition, despite the extensive knowledge and experiences of the physicians and medical specialists, it is also possible that a phenotype abnormality, which could potentially direct Phenoxome to the causal gene, might have been missed during the clinical chart review. Nonetheless, with its robust algorithm and regular database updates, Phenoxome provides an ideal platform to enable physicians and clinical researchers to interrogate the data more effectively and efficiently, in scenarios that include re-analyses.

While human factors in clinical diagnostics is difficult to model, computational tools may be deployed to account for the deficiency of HPO’s semi-automated algorithm to extract phenotype-gene relationships. For instance, a number of text mining-based tools have been developed to identify variant and gene information in biomedical literatures^48-50^. We speculate that incorporating these algorithms may serve to reinforce and supplement resources such as HPO and facilitate the clinical diagnostics by providing more supporting evidence of genes and variants of interest^51^ and therefore improve the overall effectiveness and efficiency of Phenoxome.

## Description of Supplemental Data

Supplemental Data include one figure, one table and one file.

### Acknowledgements

Clinical sequencing and analyses were supported in part by the Division of Genomic Diagnostics and the Individualized Medical Genetics Center at CHOP. The PediSeq project at CHOP was supported by NHGRI Clinical Sequencing Exploratory Research (CSER) consortium (grant U01HG006546). The authors thank Michael Hammond and Leonard Hu for their assistance in supporting the web application of this work.

## Web Resources

The URLs for data presented herein are as follows:

The Gnome Aggregation Database (gnomAD), http://gnomad.broadinstitute.org/ UCSC RefSeq, http://www.ncbi.nlm.nih.gov/RefSeq/

SnpEff, http://snpeff.sourceforge.net/

The Human Phenotype Ontology (HPO), http://human-phenotype-ontology.github.io/

Novoalign, http://www.novocraft.com/products/novoalign/

Picard, http://broadinstitute.github.io/picard/

Genome Analysis Toolkit (GATK), https://software.broadinstitute.org/gatk/

Ensembl Annotation System, http://www.ensembl.org/downloads.html

Exomiser, https://data.monarchinitiative.org/exomiser/

PhenIX, http://compbio.charite.de/PhenIX/

PediSeq Project, https://pediseq.research.chop.edu/

