## Supplementary Materials for "Rapid and Accurate Interpretation of Clinical Exomes Using Phenoxome: a Computational Phenotype-driven Approach"

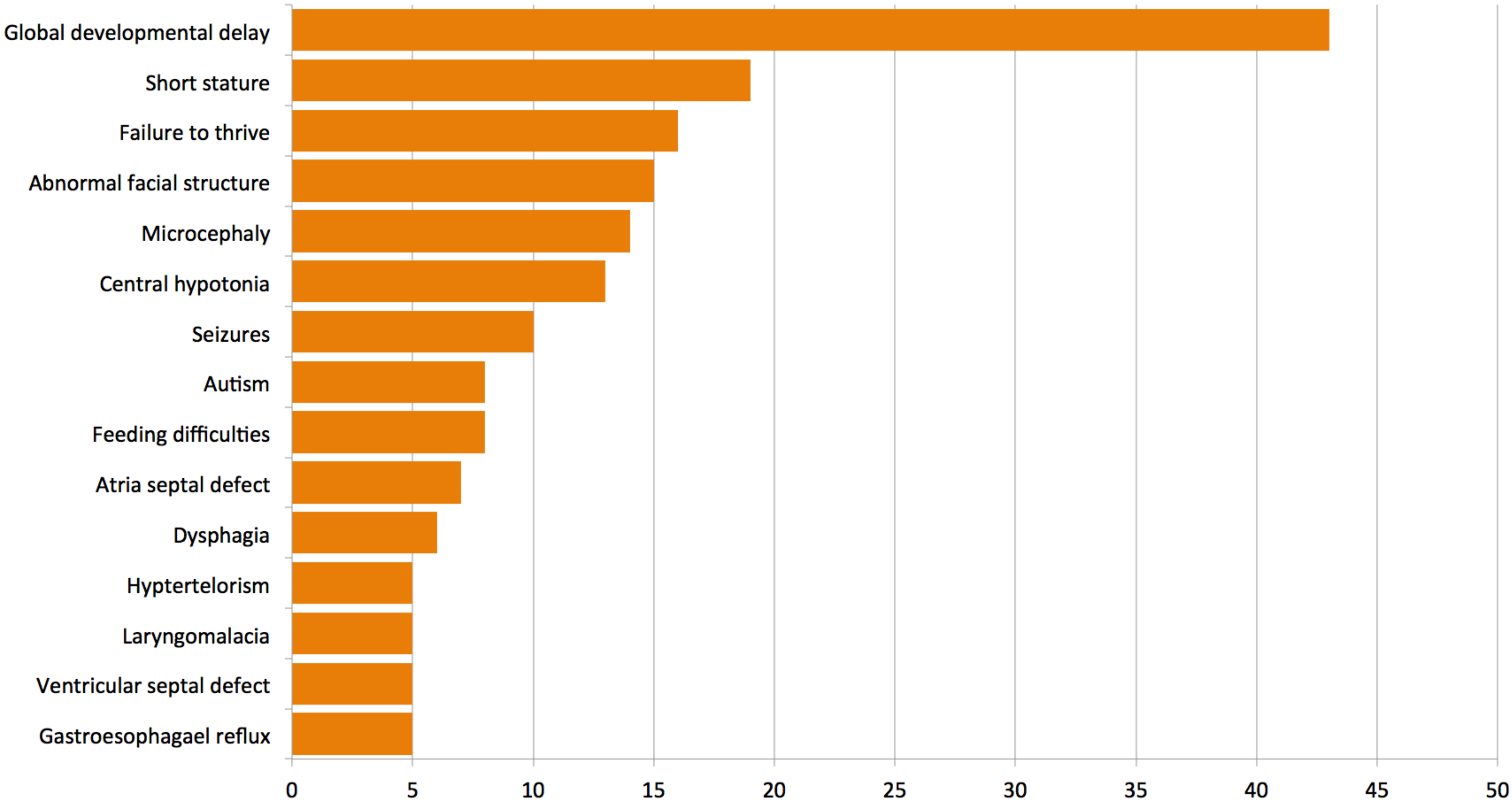


**Figure S1. Fifteen Most Frequent HPO Terms of Clinical Cohort**

*Global Development Delay* was documented for 43 patients while each of the rest terms was used to describe at least five different patients in the cohort.
