## Supplementary Materials for "Rapid and Accurate Interpretation of Clinical Exomes Using Phenoxome: a Computational Phenotype-driven Approach"

| Method | Simulation Method | # Positive Clinical Exomes |
|---|---|---|
| Masino et al. | 33 diseases with penetration | 4 |
| Phen-Gen | 765 disease from OMIM and HGMD | 16 |
| PHEVOR | 200 diseases from HGMD | 3 |
| PHIVE | 869 diseases from HGMD | 0 |
| Phenomizer | 44 diseases with penetration | 0 |
| PhenIX | 8504 simulated HGMD profiles | 52 |
| **Phenoxome** | **33 diseases with penetration** | **105** |

Table 1S. Validation Data and Resource for Computational Methods
